# Oral exposure to Perfluorooctanoic acid disrupts the microbiota-gut-liver axis and enhances the severity of chemically induced colitis in mice

**DOI:** 10.64898/2026.05.26.727994

**Authors:** Jungjae Park, Alessandra S. Miller, Gatha Pore, Maithili Banginwar, Sarah Lee, Jane Li, Eric Jung, Abigail Wagner, Jason Smith, Callan Malone, Ida Schoultz, Samira Salihovic, Colin Reardon, Mélanie G. Gareau

**Affiliations:** Department of Anatomy, Physiology, and Cell Biology, School of Veterinary Medicine, University of California, Davis, Davis, CA; School of Medical Sciences, Faculty of Medicine and Health, Örebro University, Örebro, Sweden

**Keywords:** perfluorooctanoic acid, colitis, inflammation, enterohepatic, bile acids

## Abstract

Inflammatory bowel diseases (IBD) affect millions of patients worldwide and impair quality of life. Although genetic and environmental factors are known to disrupt the gastrointestinal (GI) epithelial barrier and increase susceptibility to IBD, the precise contribution of specific environmental exposures remains unclear. Per- and polyfluoroalkyl substances (PFAS), or “forever chemicals,” are widely used in consumer products and contaminate food and water sources, resulting in chronic oral exposure worldwide. Perfluorooctanoic acid (PFOA), a common PFAS, has been epidemiologically associated with the development of IBD, particularly in older adults. Here, we assessed the effects of oral PFOA exposure on the GI tract, liver, and susceptibility to colitis. C57BL/6 mice were exposed to PFOA (0.1 mg/kg or 1.0 mg/kg) beginning at weaning (post-natal day [P]21) for a time course of 4 or 8 weeks. GI physiology/pathology (Ussing chambers; histology), expression of pro-inflammatory cytokines (qPCR), microbiota composition (16S sequencing), bile acids production (qPCR; LC/MS), and liver pathology (histology) were assessed. Colitis susceptibility was evaluated in genetically predisposed (IL10 knockout) mice, and in induced (dextran sodium sulfate [DSS]) mouse models following PFOA exposure (8 weeks at 1.0 mg/kg). Oral PFOA exposure increased intestinal permeability, mildly increased cytokine expression, altered gut microbiota composition, disrupted liver and serum bile acids, and caused hepatic hypertrophy at higher doses and longer exposure. Although PFOA did not increase disease susceptibility in genetically predisposed Il10 KO mice, it significantly worsened DSS-induced colitis, but only in male mice. Together, these findings demonstrate that early-life PFOA exposure disrupts the gut-liver axis and may contribute to colitis development in a sex dependent manner.

## INTRODUCTION

The epithelium of the gastrointestinal (GI) tract is a critical interface between the host and the external environment. As the GI tract is continuously exposed to nutrients, commensal microbes, pathogens, and environmental toxicants, the coordinated regulation of epithelial barrier function and the intestinal microbiota is essential for maintaining health. The early postnatal period is a crucial developmental window during which the intestinal barrier matures and the gut microbiota is established. Insults during this period, including stress, infection, or exposure to environmental toxicants, may disrupt the developing host-microbe interactions, leading to long-lasting health consequences.^1,2^ Inflammatory bowel diseases (IBD) are chronic, relapsing inflammatory disorders of the GI tract that affect more than 7 million people worldwide and continues to increase globally.^3–5^ IBDs are comprised of Crohn’s disease (CD) and ulcerative colitis (UC), each of which has unique features, including inflammation type and site, and origin of disease. While the precise etiology of IBD is unknown, there is strong evidence that genetic susceptibility, environmental exposures, altered microbiota, and impaired barrier function contribute to disease risk.^6,7^

Per- and polyfluoroalkyl substances (PFAS) are a class of persistent synthetic chemicals used in a variety of consumer and industrial products, including water-resistant textiles, non-stick coatings, and firefighting foams.^8^ These chemically desired properties, including stability and resistance to degradation, contribute to environmental persistence, widespread contamination, and bioaccumulation in tissues. Epidemiological studies have associated PFAS exposure, including perfluorooctanoic acid (PFOA), through contaminated drinking water with an increased risk of developing late-onset IBD.^9^ Consistent with these effects, PFOA exposure in young adult mice induced hepatotoxicity and immunotoxicity.^10^ However, whether oral PFOA exposure during early life alters GI physiology, gut-liver homeostasis, or increases susceptibility to GI inflammation remains unclear.

The gut and liver are connected via enterohepatic circulation, and bidirectional interorgan communication along the gut-liver axis helps regulate physiology.^11^ The gut microbiome is important for shaping bile acid composition and regulates host metabolic and immune responses to environmental exposures. Disruption of this microbiota-gut-liver crosstalk has been implicated in a variety of inflammatory and metabolic diseases and may represent an important mechanism by which early-life toxicant exposure modifies disease susceptibility. Here, we assessed whether oral PFOA exposure beginning at weaning disrupts GI, hepatic, and microbial homeostasis and alters susceptibility to colitis. While PFOA exposure alone did not cause overt colitis, 8 weeks of exposure impaired intestinal barrier function, altered epithelial cell proliferation, induced hepatic xenobiotic and metabolic gene transcription programs, disrupted bile acid production, and altered the gut microbiome. PFOA exposure increased the severity of dextran sodium sulfate [DSS]-induced colitis but did not enhance disease or susceptibility in genetically predisposed (interleukin-10 [IL10] knockout [KO]) mice. Together, these findings identify early-life PFOA exposure as a modifier of microbiome-gut-liver homeostasis and demonstrate that its impact on colitis severity depends on sex and inflammatory context.

## METHODS

### Experimental Design

#### Mice

Male and female C57BL/6 (Jackson Labs; bred in-house) or IL10 knockout (KO; Jackson Labs; bred in house) mice were used. PFOA exposure began at weaning (P21) and continued for 4 or 8 weeks. Tissues were collected from adult mice for analyses. Mice had *ad libitum* access to food and water during the entire experiment and were euthanized by CO2 asphyxiation followed by cervical dislocation. All procedures were approved by our institutional IACUC committee (#23455).

#### PFOA exposure

Mice were exposed to PFOA (0.1 or 1.0 mg/kg/day, Sigma Aldrich; cat #171468, M.W. 414.07) or vehicle (water) starting at weaning in the drinking water for 4 or 8 weeks, and GI and liver pathophysiology were assessed. Doses were selected based on prior studies^12^, which identified the lowest effect dose used to set the reference dose within the EPA’s drinking water lifetime health advisory level (70 parts per trillion [ppt] in 2016). Final concentration was based on the average water consumption per mouse. Body weight was recorded twice weekly.

#### Dextran sodium sulfate (DSS)

A subset of mice was administered 3% DSS (cat #160110, M.W. 36,000 – 50,00 g/mol; MP biomedicals, Irvine, California) in their drinking water for 5 days to induce colitis, followed by normal drinking water for 3 days. Mice were euthanized at day 8 post-DSS initiation. Body weight was recorded twice weekly during PFOA exposure and daily during DSS-induced colitis.

### Ussing Chambers

Ussing chamber studies were performed as described previously.^13,14^ Distal ileum and colon segments were collected immediately after euthanasia and placed in cold carbogenated Ringer’s buffer (pH 7.35 ± 0.02). Tissue segments (1 cm) were opened longitudinally, rinsed gently to remove luminal contents, and mounted. Tissue was then mounted in Ussing chambers (Physiological Instruments, Reno, NV), and baseline ion transport (short-circuit current [Isc]) and conductance (G) were measured to assess ion transport and tight junction permeability, respectively. FITC-labeled dextran (4kDa) was added to the mucosal buffer, and transport to the serosal side was assessed by sampling every 30 minutes for 2h. Fluorescence was measured using a plate reader at excitation/emission wavelengths of [490/520] nm, and flux was calculated using a standard curve (µg/cm²/hour).

### Targeted LC-MS/MS Analysis

Serum and liver samples were collected from mice exposed to PFOA (1mg/Kg, 8 weeks) or vehicle for 8 weeks and stored at −80°C until analysis. Targeted metabolomic profiling was performed to quantify PFAS and bile acids in serum and liver extracts using liquid chromatography-mass spectrometry (Acquity PREMIER UHPLC-Xevo TQXS MS/MS, Waters, MA, USA). Liver samples were homogenized prior to extraction. An isotope-labelled internal standards mixture comprising 26 individual compounds (Tauroursodeoxycholic acid-d4, Ursodeoxycholic acid-d4, Taurocholic acid-d4, Glycocholic acid-d4, Taurochenodeoxycholic acid-d9, Glycochenodeoxycholic acid-d4, Chenodeoxycholic acid-d9, Glycodeoxycholic acid-d4, Deoxycholic acid-d4, Taurolithocholic acid-d4, Glycolithocholic acid-d4, Lithocholic acid-d4, Cholic acid-d4, ¹³C₄-PFBA, ¹³C₅-PFPeA, ¹³C₅-PFHxA, ¹³C₄-PFHpA, ¹³C₈-PFOA, ¹³C₉-PFNA, ¹³C₆-PFDA, ¹³C₇-PFUnDA, ¹³C₂-PFDoA, ¹³C₂-PFTeDA, ¹³C₃-PFBS, ¹³C₃-PFHxS, ¹³C₈-PFOS) was added to both the serum and liver homogenate samples and metabolites were isolated using a previously developed sample preparation and extraction method.^15^ Extracted samples were further clarified by centrifugation, transferred to LC-MS vials, and analyzed using a targeted PFAS and bile acid panel. PFAS and Bile acid species were separated using an Acquity Premier BEH C18 analytical column and detected by MS using optimized MRM transitions for individual PFAS and bile acids. Quantification was performed using a fetal bovine serum matrix matched calibration curve 0.5-1000ng/mL spiked with 39 individual PFAS and bile acids. The quantified metabolites included 21 individual PFAS congeners and 18 primary, secondary, conjugated, and unconjugated bile acids. Absolute bile acid concentrations were compared between vehicle- and PFOA-exposed mice to assess the effects of PFOA exposure on systemic and hepatic bile acid metabolism. Data were normalized to sample volume for serum and tissue weight or protein content for liver samples, as appropriate.

### Quantitative PCR

Tissues (colon, ileum, and liver) were collected in TRIzol (Invitrogen, Carlsbad, California) and frozen at -80°C. RNA was extracted according to the manufacturer’s instructions, and RNA concentration and purity were assessed using a NanoDrop. Synthesis of cDNA was performed using the iScript reverse transcriptase kit (Bio-Rad, Hercules, California). Quantitative PCR was performed using SYBR Green chemistry on a QuantStudio 6 machine. Relative gene expression was calculated using the ΔΔCt method and normalized to [housekeeping gene(s)]. Primer sequences are listed in Table 1.

**Table 1.**
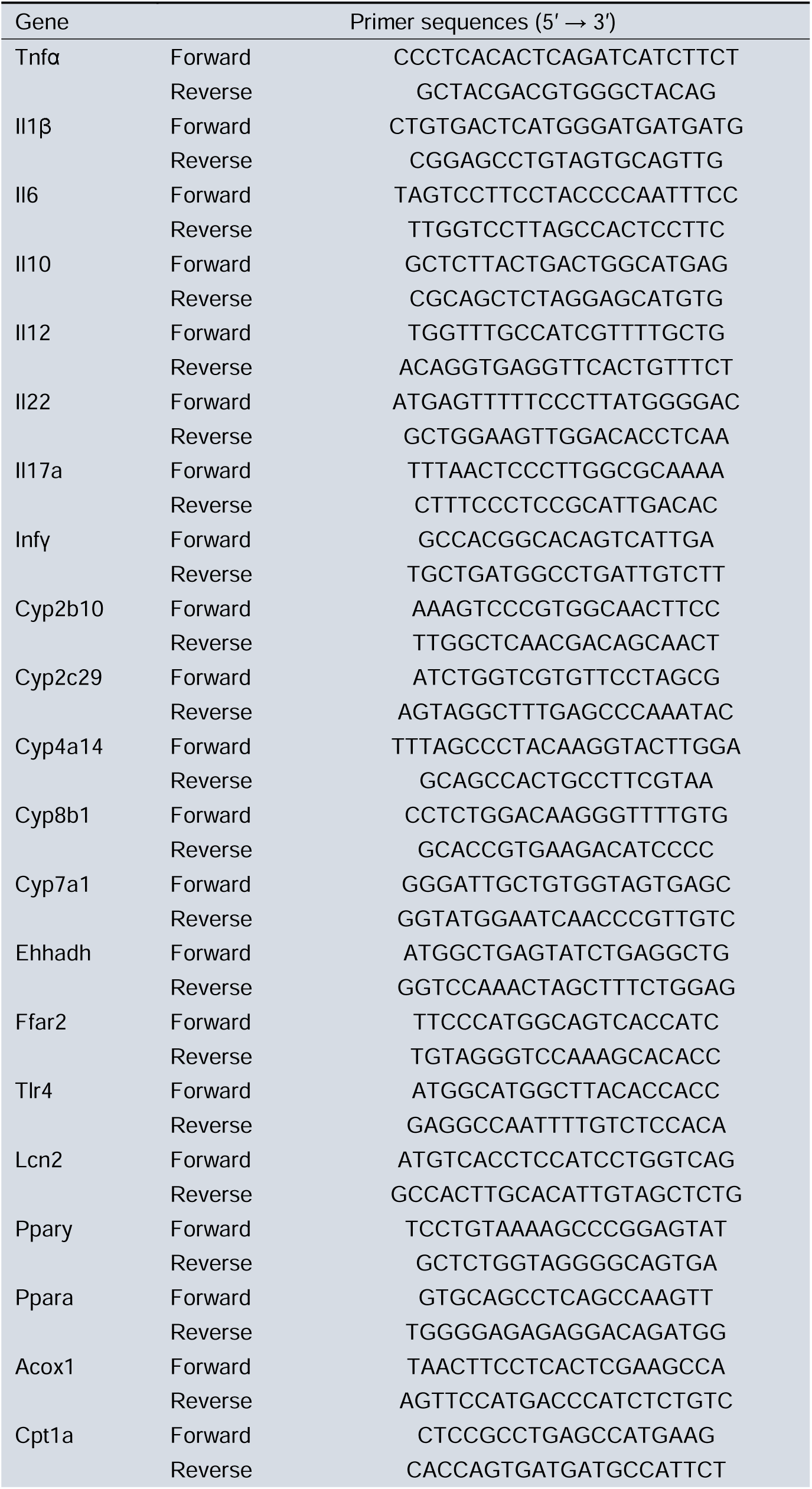
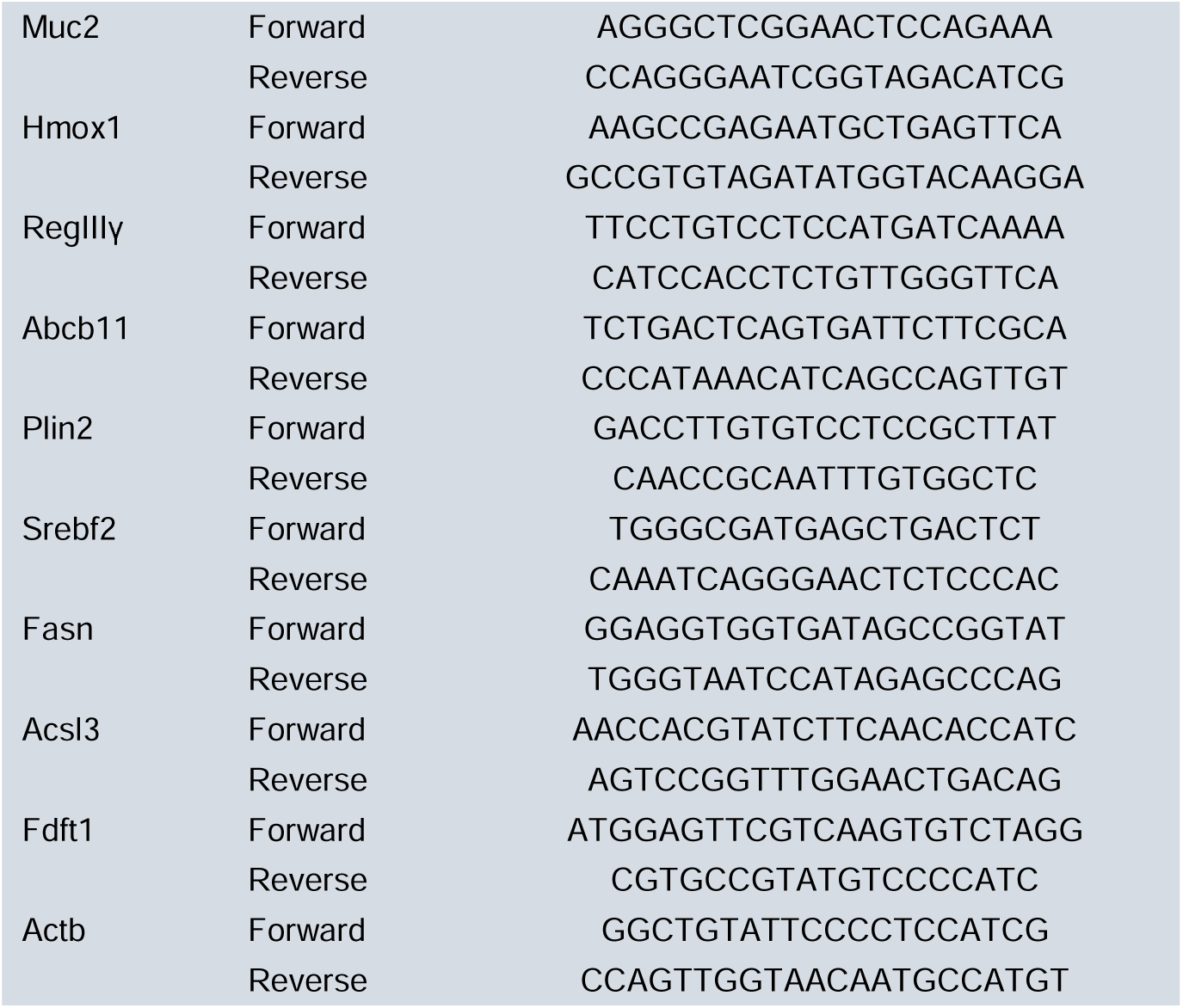
Primer sequences.

### Histology

#### Hematoxylin and Eosin (H&E)

Ileum, colon, and liver were fixed in 10% formalin for a minimum of 48 hours, transferred to 70% ethanol, processed for paraffin embedding, and sectioned using a microtome (5 µm). Slides were then deparaffinized with Histoclear, rehydrated, stained with H&E, and imaged by light microscopy (Nikon Eclipse E600 using Nikon DS-U3 digital sight). Colonic and ileal crypt depth measurements were quantified from 10 well-oriented crypts/villi per mouse (FIJI/ImageJ). Hepatic cellularity was quantified by counting nuclei in 2 randomly selected fields per mouse at 20X (FIJI/ImageJ). DSS-induced colitis severity was assessed using a composite score that included epithelial injury, architectural distortion, inflammatory cell infiltration, and ulceration.^16^ All imaging and associated quantification were obtained using coded slides generated by a random number generator and averaged for each animal.

#### Periodic Acid Schiff (PAS)

Goblet cells were quantified using periodic acid-Schiff (PAS) staining in colonic tissues. Slides were deparaffinized with Histoclear, rehydrated, and incubated in 1% periodic acid for 5 minutes. Sections were then incubated in Schiff’s reagent for 15 minutes, washed in running tap water, counterstained with hematoxylin for 1 minute, and imaged by light microscopy. PAS-positive goblet cells were quantified using FIJI/ImageJ across two to three representative sections per mouse and averaged. Data were expressed as total PAS-positive cells per animal.^17^

### Immunofluorescence

Epithelial cell proliferation was assessed by Ki67 immunostaining using formalin-fixed, paraffin-embedded colon tissue. Using a mouse-on-mouse immunodetection kit (Vector Labs), sections were stained with a primary rabbit anti-Ki67 antibody (LS-C141898; Lifespan Biosciences) followed by Alexa Fluor 555 goat anti-rabbit [A21428; Life Technologies] secondary antibody and co-staining with the IEC marker E-Cadherin (CDH1) using mouse anti-CDH1 (CM1681; ECM Biosciences) and streptavidin-Alexa Fluor 647 (S32357; Invitrogen). Nuclei were stained with 4′,6-diamidino-2-phenylindole (DAPI). Images were acquired on a Leica SP8 confocal microscope (40x objective, 0.7 µm step size), and overlapping tiles were stitched and Huygens deconvolved as needed, followed by quantification of Ki67^+^CDH1^+^ cells/crypt in Imaris 8.2 software.

### Microbiome 16S rRNA sequencing

Fecal pellets were collected at euthanasia, and bacterial DNA was extracted from feces (∼100 mg) using the DNeasy PowerSoil Pro Kit (Qiagen, Irvine, CA) per the manufacturer’s instructions, followed by library preparation. Bacterial 16S rRNA genes were amplified by PCR using a primer set that targets the V3-V4 region (515 F: 5’-XXXXXXXXGTGTGCCAGCMGCCGCGG TAA-3’; 806 R: 5’-GGACTACHVGGGTWTCTAAT-3’), where “X” denotes an 8-bp sample-specific barcode and “GT” represents the Illumina adapter sequence. Extraction and PCR negative controls, including ultrapure water and DNA elution buffer, were included to monitor reagent and environmental contamination. Libraries were submitted to the UC Davis Genome Center and sequenced using paired-end 250-bp reads on the Illumina MiSeq platform. Microbiome analysis was performed using QIIME 2^18^ and R, following a previously described workflow.^19^ Raw reads were demultiplexed using Sabre and imported into QIIME2. Reads were quality filtered and denoised using DADA2^20^, including primer trimming, read truncation at 240 nucleotides, paired-end merging, and chimera removal. Taxonomy was assigned using q2-feature-classifier and the SILVA 138.2 database trained for the amplified region.^21^ ASV tables and phylogenetic trees were exported to R for analysis with phyloseq.

Alpha diversity, beta diversity, and taxonomic relative abundance were calculated and visualized in R using phyloseq and ggplot2. All computational analyses were conducted using the Farm Cluster, a Linux-based high-performance computing system at the College of Agricultural and Environmental Sciences, University of California, Davis. The 16S rRNA sequencing data have been deposited in the NCBI Sequence Read Archive under BioProject ID PRJNA1470048.

### Statistical Analysis

Statistical analyses were performed using GraphPad Prism version 10 (GraphPad Software, San Diego, CA). Power calculations were performed using G*power (α = 0.05, [1-β] = 0.8) to determine sample size. Results were analyzed by 1- or 2-way ANOVA with post-hoc testing (i.e. Tukey’s or Bonferroni) as appropriate. Normality was assessed by the Shapiro-Wilk test and parametric or non-parametric testing was performed as appropriate. Data is presented as mean ± standard error of mean (SEM) and n represents individual mice unless otherwise stated. Significance was defined as *P<0.05, **P<0.01, ***P<0.001, or ****P<0.0001.

For the microbiome analysis, statistical analyses were performed in R (version 4.5.3). Data distribution and variance assumptions were evaluated prior to analysis, with normality assessed using the Shapiro-Wilk test and homogeneity of variance tested using Bartlett’s test for α-diversity and relative abundance datasets. Since the data did not meet parametric assumptions, nonparametric statistical approaches were applied. α-Diversity indices and relative abundances of bacterial taxa were analyzed using the Kruskal-Wallis rank-sum test, followed by post hoc multiple comparisons using Dunn’s test or Conover’s test, implemented via the agricolae package in R. β-Diversity was assessed using Bray–Curtis dissimilarity metrics, which were visualized by principal coordinates analysis (PCoA). Differences in community structure between groups were statistically evaluated using permutational multivariate analysis of variance (PERMANOVA) with 999 permutations, implemented in the vegan package. To account for multiple comparisons in taxa-level analyses, P values were adjusted using the Benjamini–Hochberg false discovery rate (FDR) correction. Statistical significance was defined as an adjusted P value (Padj) < 0.05, with 0.05 ≤ Padj < 0.10 considered a trend toward significance.

## RESULTS

### PFOA exposure disrupts intestinal barrier function and induces hepatic metabolic remodeling

The effects of dose and duration of oral PFOA exposure on the GI tract and liver were assessed by exposing mice to low (0.1 mg/Kg) or high (1 mg/Kg) PFOA doses in drinking water beginning at weaning for 4 or 8 weeks (**Fig 1A**). Body weight was monitored before and throughout PFOA exposure, revealing no significant differences in post-weaning growth versus vehicle control-treated mice (**Supplemental Fig 1A**). GI physiology was assessed in the colon and ileum by Ussing chambers. Colonic FITC-dextran permeability increased after exposure to low- and high-dose of PFOA, although this effect did not follow a linear dose response (**Fig 1B**). In contrast, ileal FITC-dextran permeability was elevated only in mice exposed to low, but not high-dose PFOA (**Fig 1B**). Although colonic and ileal basal secretory state and conductance were not altered at either dose, 8-week PFOA exposure significantly reduced carbachol-evoked Isc responses in the colon, but not in the ileum (**Supplementary Fig 1B**).

**Figure 1.**
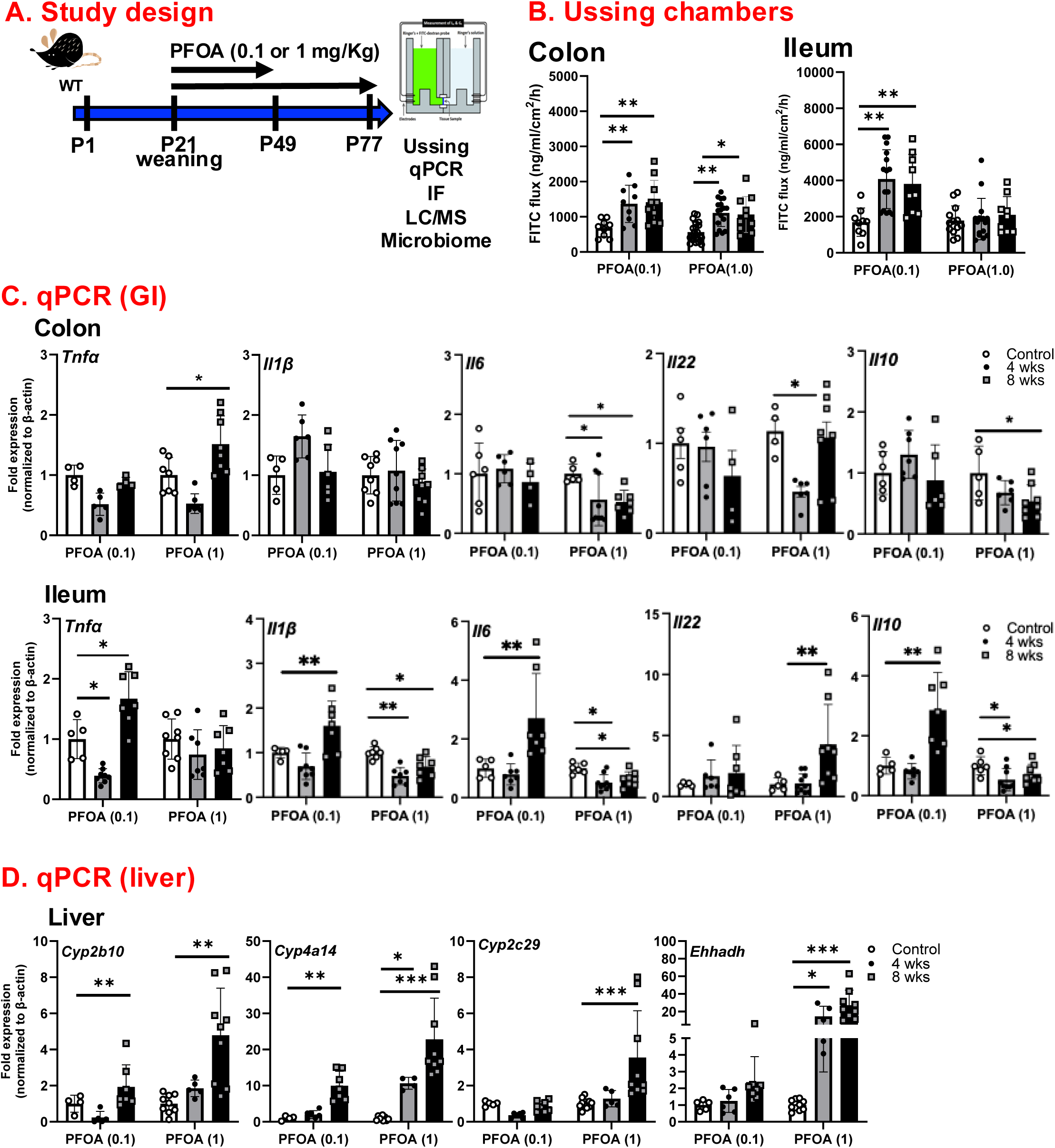
Chronic PFOA exposure increases colonic permeability and inflammatory gene expression. Experimental design schematic for dose (0.1 – 1 mg/kg) and duration (4 and 8 weeks) of PFOA exposure (**A**). Permeability assessed by flux of FITC-labeled dextran in Ussing chambers of distal colon and distal ileum from vehicle control and PFOA-exposed mice (**B**). Inflammation was assessed by qPCR in the colon and ileum (*Tnfa, Il1b, Il6, Il22, Il10*) and metabolism/detoxification (*Cyp2b10, Cyp4a14, Cyp2c29*) and peroxisomal enzymes (*Ehhadh*) were assessed in the liver (**C**). Data are presented as mean ± SEM; n=4-8 [qPCR, 8-20 [Ussing]), *p<0.05, **p < 0.01, ***p<0.001 by two-way ANOVA.

Given these PFOA exposure-induced changes in GI permeability, inflammation was assessed by qPCR of cytokine and host-protective gene expression. In the colon, low-dose PFOA exposure did not significantly alter the expression of inflammatory cytokine genes. In contrast, high-dose PFOA exposure increased *Tnf*α expression and reduced Il6 and Il10 expression after 8 weeks (**Fig 1C**). Il22 expression decreased only in the high-dose 4-week exposed group, whereas *Il1*β expression remained unchanged across doses and timepoints (**Fig 1C**). In the ileum, low-dose PFOA exposure increased the expression of *Tnf*α, *Il1*β, *Il6*, and *Il10* after 8 weeks (**Fig 1C**). In contrast, high-dose PFOA exposure suppressed *Il1*β, *Il6*, and *Il10* expression at both 4 and 8 weeks without altering *Tnf*α expression, whereas *Il22* expression increased only after 8 weeks (**Fig 1C**). In the liver, we evaluated expression of genes associated with xenobiotic responses, peroxisome proliferator–activated receptor alpha (Pparlll) regulated lipid metabolism, and bile acid homeostasis. PFOA exposure increased hepatic *Cyp2b10 and Cyp4a14* expression at both doses, whereas *Cyp2c29* and *Ehhadh* expression increased only in mice exposed to high-dose PFOA (**Fig 1D**). Together, these data indicate that oral PFOA exposure beginning at weaning reduces barrier function and induces hepatic metabolic response genes, with effects that vary by tissue and dose but not as a linear dose-response.

Given evidence of PFOA exposure-induced changes in intestinal barrier function and hepatic gene expression, histology was performed to characterize tissue pathology. PFOA exposure (1 mg/Kg for 8 weeks) significantly reduced colonic crypt depth (**Fig 2A**), whereas ileal histomorphology was unaltered (**Suppl Fig 1C**). This reduced colonic crypt depth was accompanied by a significant reduction in the number of Ki67+ cells, suggesting that altered epithelial cell dynamics were driven by a reduction in proliferating cells (**Fig 2B**). In contrast, PAS staining for goblet cells revealed no change in colonic goblet cell abundance (**Fig 2C**). In the liver, a decrease in the number of hepatocytes was found in PFOA-exposed mice (**Fig 2D**), which was coupled with an increase in the liver-to-body ratio (**Fig 2E**).

**Figure 2.**
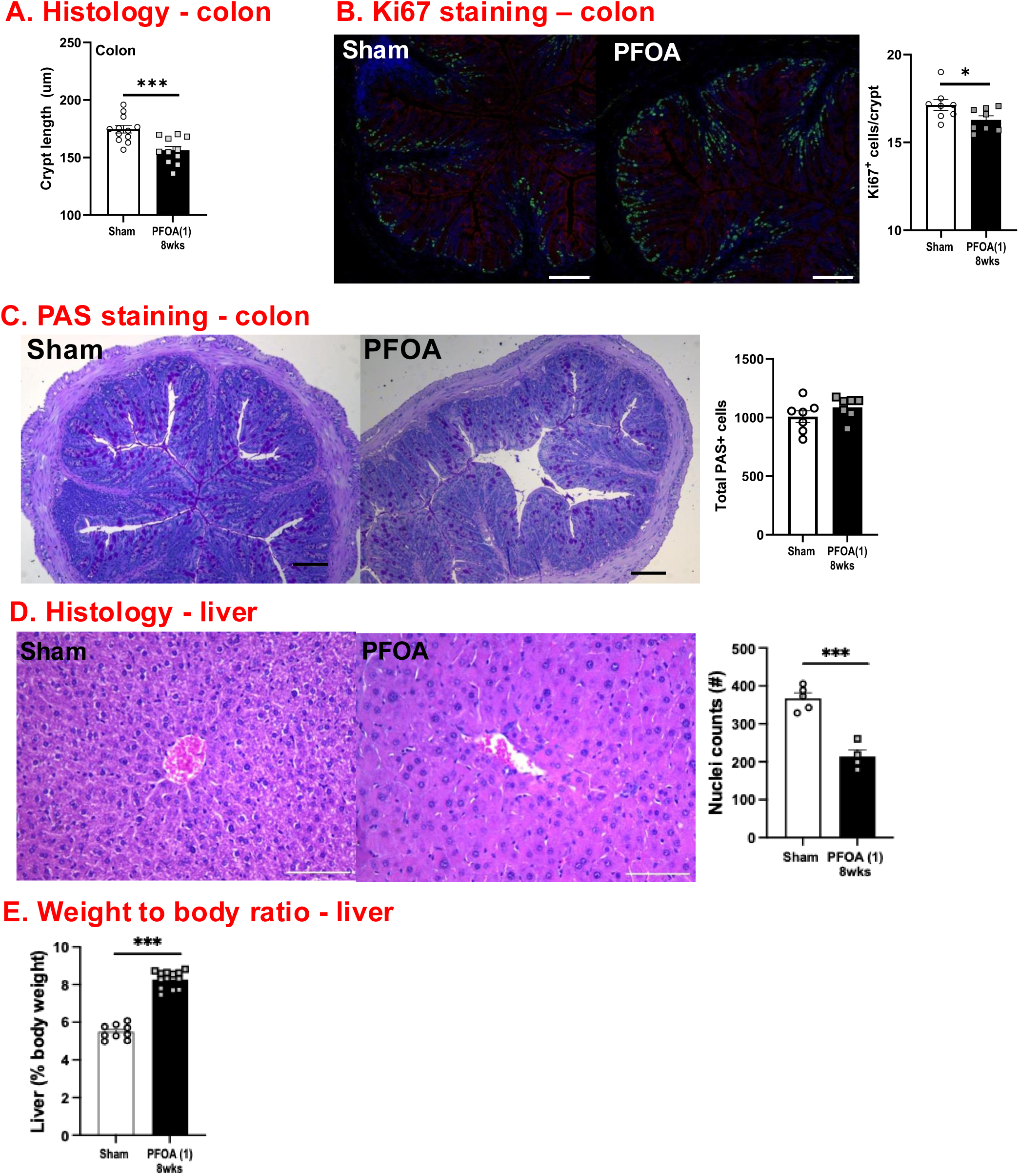
Chronic PFOA exposure induces colonic and hepatic histopathology. Histological scoring of the colon and liver following PFOA exposure (1mg/kg, 8 weeks). Crypt depth was quantified (**A**). Representative immunofluorescence images of colon sections stained for Ki67 [green], CDH1/E-cadherin [red] nuclei DAPI [blue], with quantification of Ki67+ IEC per crypt (**B**). Representative periodic acid–Schiff (PAS) staining and quantification of total goblet cell-associated mucins (magenta) (**C**). Representative hematoxylin and eosin (H&E) staining and nuclei counts of hepatocytes in the liver (**D**) Liver-to-body weight ratio following 8 weeks of PFOA exposure (**E**). Data are presented as mean ± SEM; n=7-12; *p<0.05, **p < 0.01, ***p<0.001 by two-way ANOVA with Bonferroni correction, (Scale bar = 100 μm).

### PFOA exposure alters primary and secondary bile acids in both serum and liver

Given the role of bile acids as central mediators of the gut-liver axis, bile acid concentrations were measured in serum and liver by LC-MS/MS. In serum, cholic acid (CA) was increased, whereas glycolithocholic acid (GLCA), taurodeoxycholic acid (TDCA), and tauroursodeoxycholic acid (TUDCA) were decreased in PFOA-exposed mice compared with vehicle controls (**Fig 3A**). In the liver, glycocholic acid (GCA) was significantly increased whereas taurochenodeoxycholic (TCDCA), TDCA, taurohyodeoxycholic acid (THDCA), and TUDCA were reduced. Together, these findings suggest that PFOA exposure causes tissue-specific changes in bile acid composition.

**Figure 3.**
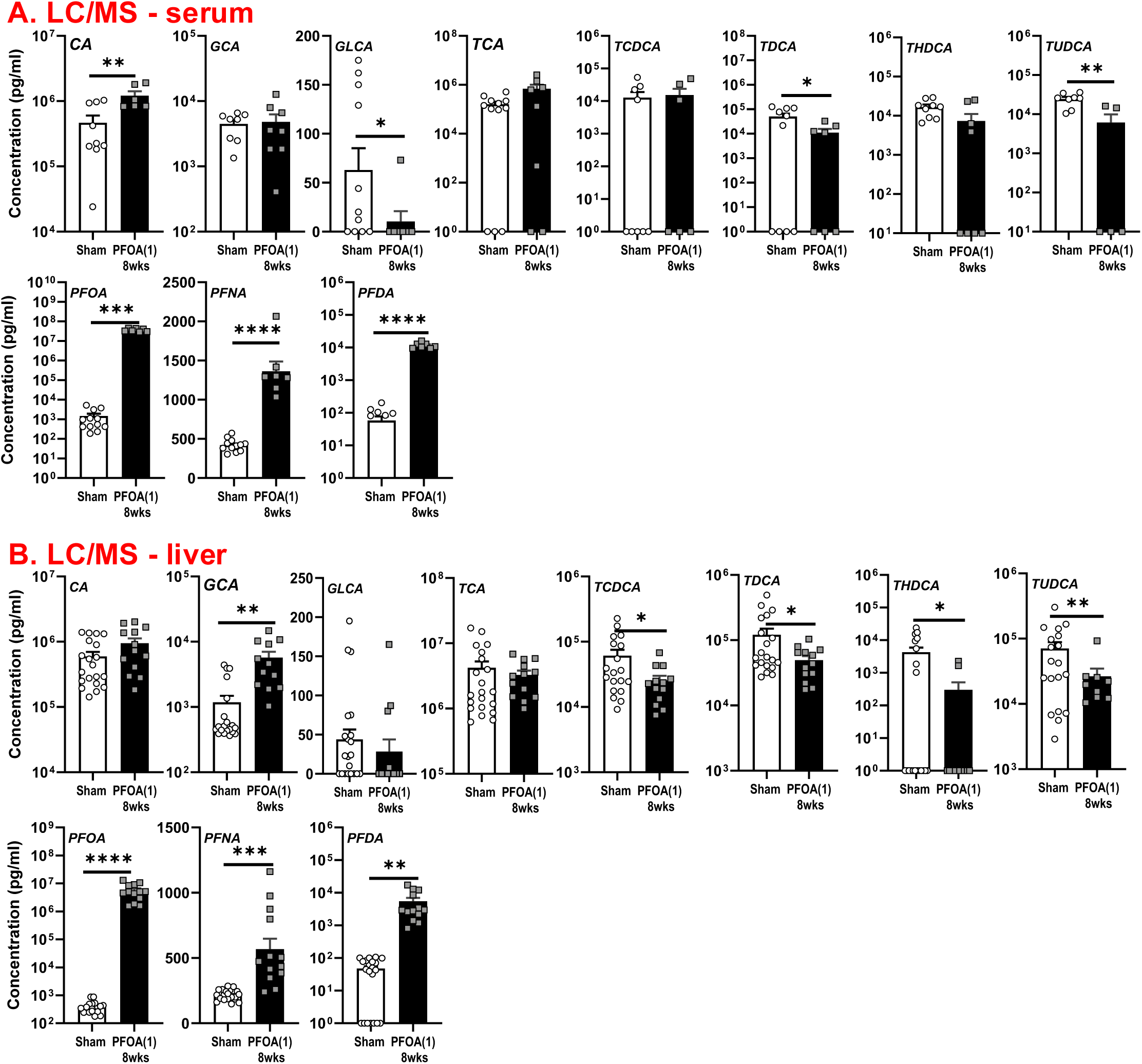
Chronic PFOA exposure alters serum and hepatic bile acid concentration. Targeted LC-MS/MS analysis of serum bile acids following PFOA exposure (1mg/kg, 8 weeks) (A). Targeted LC-MS/MS analysis of hepatic bile acids following chronic PFOA exposure (B). Individual bile acid species quantified included cholic acid (CA), glycocholic acid (GCA), glycolithocholic acid (GLCA), taurocholic acid (TCA), taurochenodeoxycholic acid (TCDCA), taurodeoxycholic acid (TDCA), taurohyodeoxycholic acid (THDCA), and tauroursodeoxycholic acid (TUDCA). Data are presented as mean ± SEM; n=10-18, *p<0.05, **p < 0.01, ****p<0.001 by Student’s T-test.

We further quantified PFAS concentrations in serum and liver to confirm systemic and hepatic accumulation following oral exposure. As expected, PFOA was detected at high levels in both serum and liver, with additional PFAS, including perfluorononanoic acid (PFNA) and perfluorodecanoic acid (PFDA), also identified in exposed animals, albeit at much lower concentrations. These concentrations are higher than those typically reported in broad epidemiological surveys, but are consistent with exposure levels observed in populations living near PFAS production facilities and among workers at PFAS-manufacturing sites.^22^

### PFOA exposure reshapes the gut microbiome composition

Since PFAS exposure is associated with disruption of infant gut microbiome colonization^23^, we assessed whether oral PFOA exposure at weaning altered the microbiome using 16S rRNA gene sequencing of fecal samples. Exposure to PFOA increased alpha diversity of the microbiome, as measured by Shannon and Chao1 indices, compared with vehicle-exposed controls (**Fig 4A**). PCoA was used to assess beta diversity, with data showing that PFOA-exposed mice clustered separately from vehicle-exposed mice, indicating that their microbial community structure was altered by exposure (**Fig 4B**). To further elucidate the impact of PFOA exposure on microbiota composition, relative abundances across multiple bacterial taxonomic levels, including phylum, family, and genus, were determined. At the phylum level, PFOA exposure increased the relative abundance of *Firmicutes*, *Actinobacteriota*, and *Proteobacteria* and decreased *Bacteroidota* and *Verrucomicrobiota* compared to controls (**Fig 4C**). Family-level analysis revealed broad remodeling of *Firmicutes*-associated taxa, including expansion of *Enterococcaceae* and *Streptococcaceae* and reduction of Lachnospiraceae and *Ruminococcaceae*, two families that include prominent short-chain fatty acid (SCFA)-producing taxa (**Supplementary Fig. 2**). At the genus level, PFOA exposure increased *Bifidobacterium*, *Blautia*, *Escherichia-Shigella*, *Peptoclostridium*, and *Streptococcus,* while decreasing *Akkermansia*, *Lachnospiraceae_NK4A136_*group, *Lactobacillus*, *Muribaculaceae*, and *Turicibacter* (**Fig 4C**). These results demonstrate that PFOA exposure selectively alters specific microbial populations, including taxa associated with SCFA production and mucosal barrier regulation.

**Figure 4.**
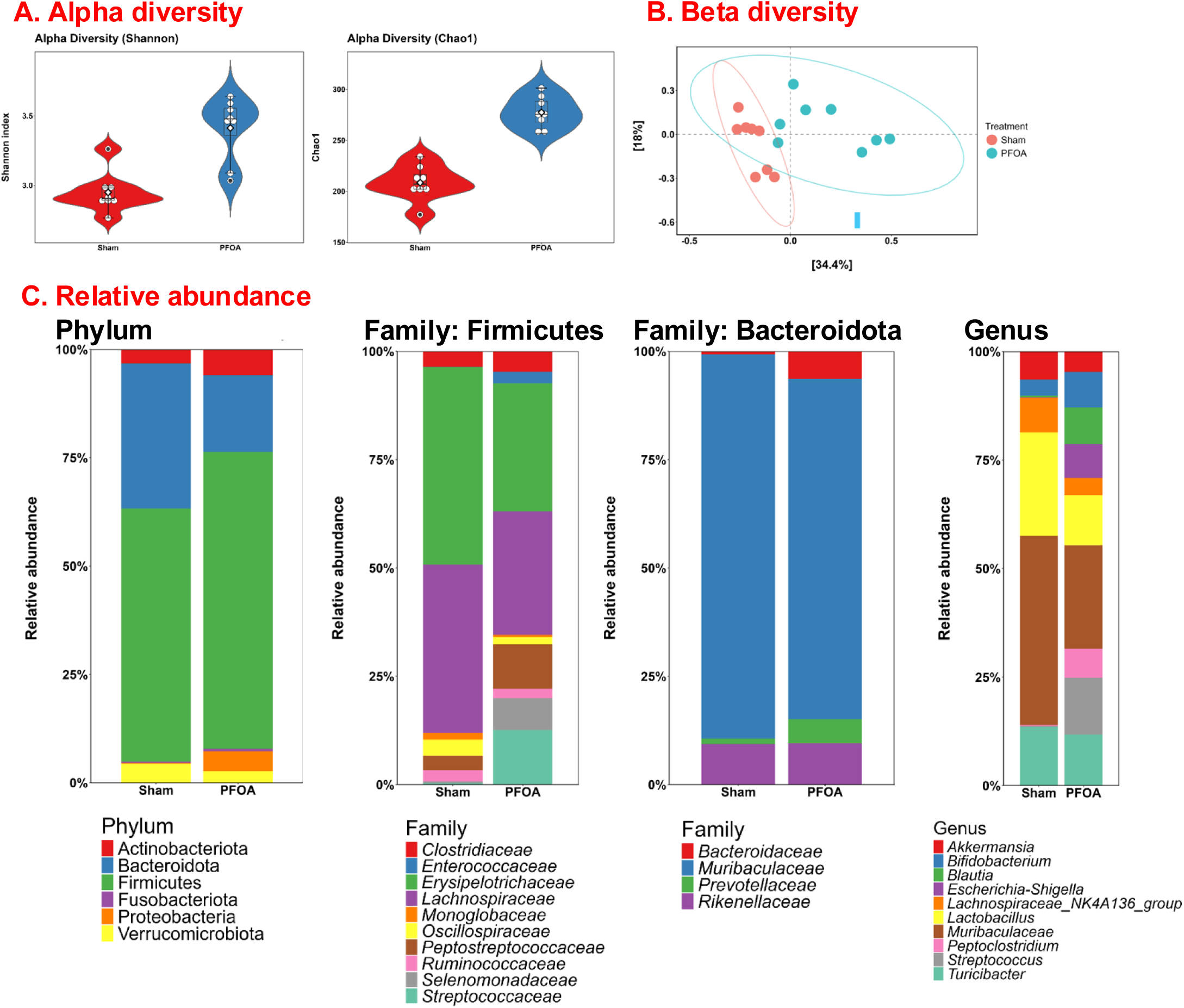
Chronic PFOA exposure impacts gut microbial diversity and taxonomic composition. Fecal microbiota via 16S sequencing of PFOA exposed (1mg/Kg, 8 weeks) vs sham-exposed mice. Alpha diversity was assessed using the Shannon diversity and Chao1 richness indices (**A**). Beta diversity analysis of microbial community composition using principal coordinate analysis (PCoA) based on Bray-Curtis dissimilarity metrics (**B**). Relative abundance analyses of fecal microbiota at the phylum, family, and genus levels following chronic PFOA exposure (**C**). n=8 mice per group.

### PFOA exposure does not worsen GI physiology in Il10-deficient mice despite altered inflammatory gene expression

Given the increased GI permeability and altered cytokine gene expression following PFOA exposure, susceptibility to GI inflammation was assessed in a genetic model of colitis. IL10^-/-^mice, which spontaneously develop colitis by approximately 3-4 months of age in a microbiome-dependent manner^24^, were exposed to PFOA (1mg/kg for 8 weeks) starting at weaning (**Fig 5A**). PFOA exposure caused a modest reduction in weight gain in male IL10^-/-^ mice, whereas female mice showed a modest increase in weight gain compared to vehicle controls (**Fig 5B**). Colonic physiology was not altered between vehicle- and PFOA-exposed IL10^-/-^ mice as determined by Ussing chambers (**Fig 5C**). However, PFOA exposure increased colonic expression of *Tnf*α*, Il6, Il17a,* and *Ifn*γ expression coupled with a reduction in *Il22* (**Fig 5D**). In the liver, similar to WT mice, PFOA exposure increased expression of metabolic and xenobiotic-response genes in IL10^-/-^ mice (**Fig 5D**). Together, these findings indicate that PFOA exposure modestly alters colonic inflammatory gene expression in IL10^-/-^ mice, which was not accompanied by worsening of GI physiology or overt disease severity.

**Figure 5.**
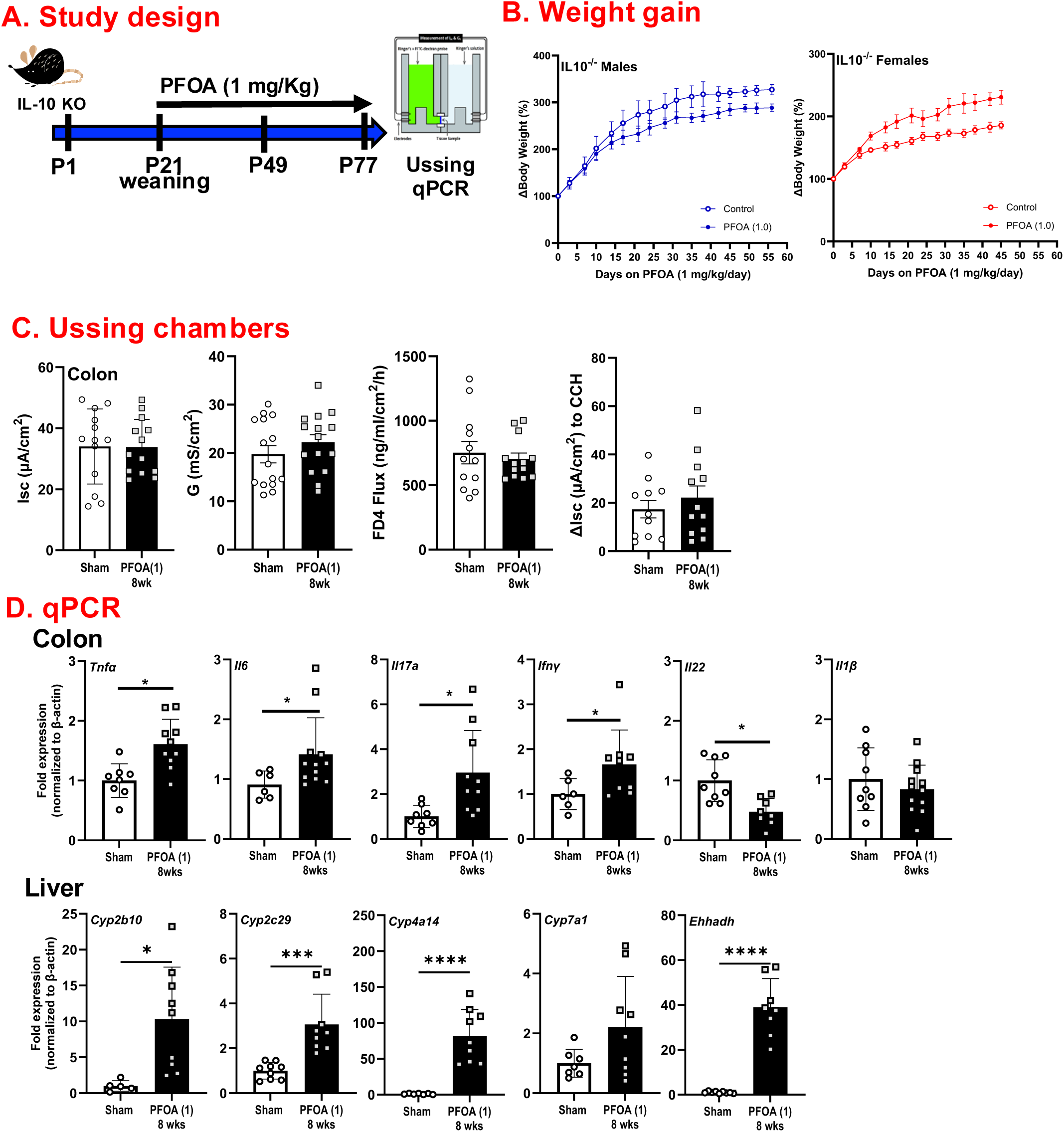
Chronic PFOA exposure does not impact colitis in an IL10 knockout model. Experimental design schematic of PFOA exposure (1mg/kg, 8 weeks) in IL10^-/-^ mice. Body weight was monitored throughout exposure (**A**). GI physiology was assessed for ion secretion (short-circuit current [Isc], conductance [G], flux of FITC dextran, and carbachol induced Isc) by Ussing chambers (**B**). Colonic inflammation (*Tnfa, Il6, Il17a, Ifng, Il22, Il1b*) and metabolism/detoxification (*Cyp2b10, Cyp4a14, Cyp2c29*) and peroxisomal enzymes (*Ehhadh*) in the liver were analyzed by qPCR (**D**). Data are presented as mean ± SEM; n=8-16, *p<0.05, **p < 0.01, ****p<0.001 by Student’s T-test.

### DSS-induced colitis is exacerbated in male mice exposed to PFOA

As PFOA exposure did not increase disease susceptibility in a genetic mouse model of colitis, a chemically-induced model of colonic inflammation was used, and disease severity was assessed following PFOA exposure. DSS-induced colitis was initiated after exposure to PFOA (1.0 mg/kg, 8 weeks) beginning at weaning, and disease severity was assessed at 8 days post-DSS (**Fig 6A**). As expected, DSS induced significant weight loss and colitis in all groups, coupled with decreased colon length compared to sham controls (**Fig 6B**). Histological analysis confirmed overt damage in DSS-induced colitis, characterized by immune cell infiltration and crypt disruption (**Fig 6C**). Despite similar effects on colon length and body weight, histological disease scoring revealed that prior PFOA exposure significantly worsened DSS-induced disease severity in only male mice (**Fig 6C)**. GI physiology via Ussing chambers indicated that PFOA enhanced conductance and FITC-dextran flux in male mice subjected to DSS-induced colitis. In female mice, prior PFOA exposure increased FITC-dextran flux, and PFOA alone reduced Isc and G (**Fig 6D**).

**Figure 6.**
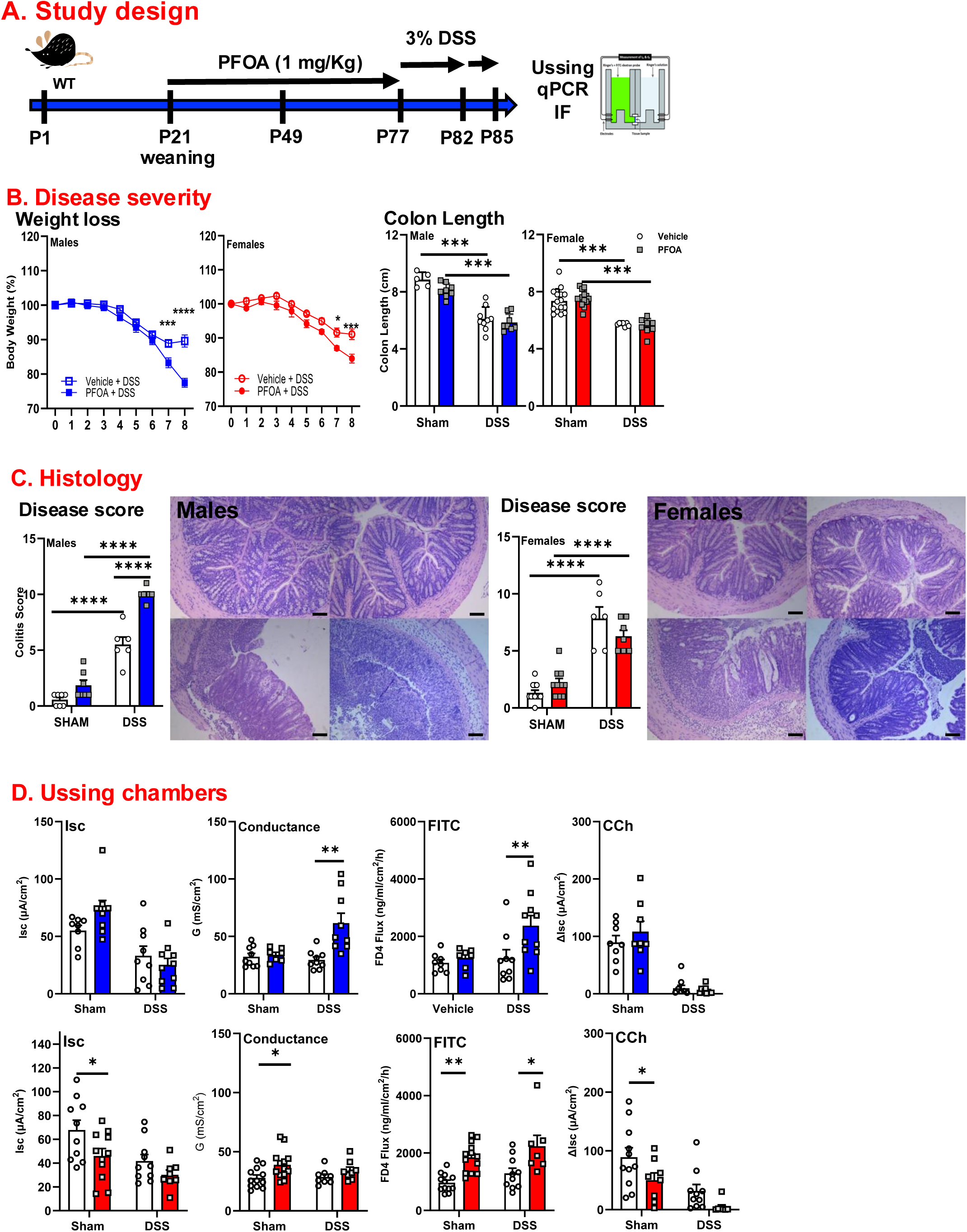
Chronic PFOA exposure exacerbates DSS-induced colitis severity in a sex-dependent manner. Experimental design schematic of DSS-induced colitis plus PFOA exposure (1mg/Kg, 8 weeks) (**A**). Body weight loss was monitored during DSS-induced colitis, and colon length was determined at day 8 post-DSS (**B**). Disease severity was quantified by histological scoring (**C**; representative images). Colonic pathophysiology was assessed using Ussing chambers, including short-circuit current [Isc], conductance [G], FITC-labeled dextran flux, and stimulated ion transport with carbachol) (**D**). Data are presented as mean ± SEM; n=8-16, *p<0.05, **p < 0.01, ***p<0.001 by two-way ANOVA.

The enhanced DSS-induced histological disease severity observed in male mice was further assessed by inflammatory gene expression. Colonic inflammatory and host-protective gene expression was measured by qPCR. In male mice, prior PFOA exposure enhanced DSS-induced colonic expression of multiple inflammatory genes including *Tnf*α, *Il1*β, *Il6*, *Ifn*γ, and also increased expression of *Il10 and Il22* (**Fig 7A** and **Supplemental Fig 3**). Colonic gene expression further supported the idea that prior PFOA exposure altered the epithelial response to DSS-induced injury. PFOA exposure alone reduced Muc2 expression, which was further amplified by DSS-induced colitis, suggesting impaired mucus barrier-associated responses during inflammatory injury (**Fig 7A**). In contrast, DSS-induced expression of the antimicrobial lectin RegIIIg was attenuated by prior PFOA exposure, whereas Lcn2 was markedly elevated only in PFOA-exposed DSS-treated mice (**Fig 7A**). Together, these changes suggest that prior PFOA exposure does not simply amplify epithelial defense programs but rather shifts the colonic response toward an injury-associated inflammatory state with impaired mucus and antimicrobial barrier features.

**Figure 7.**
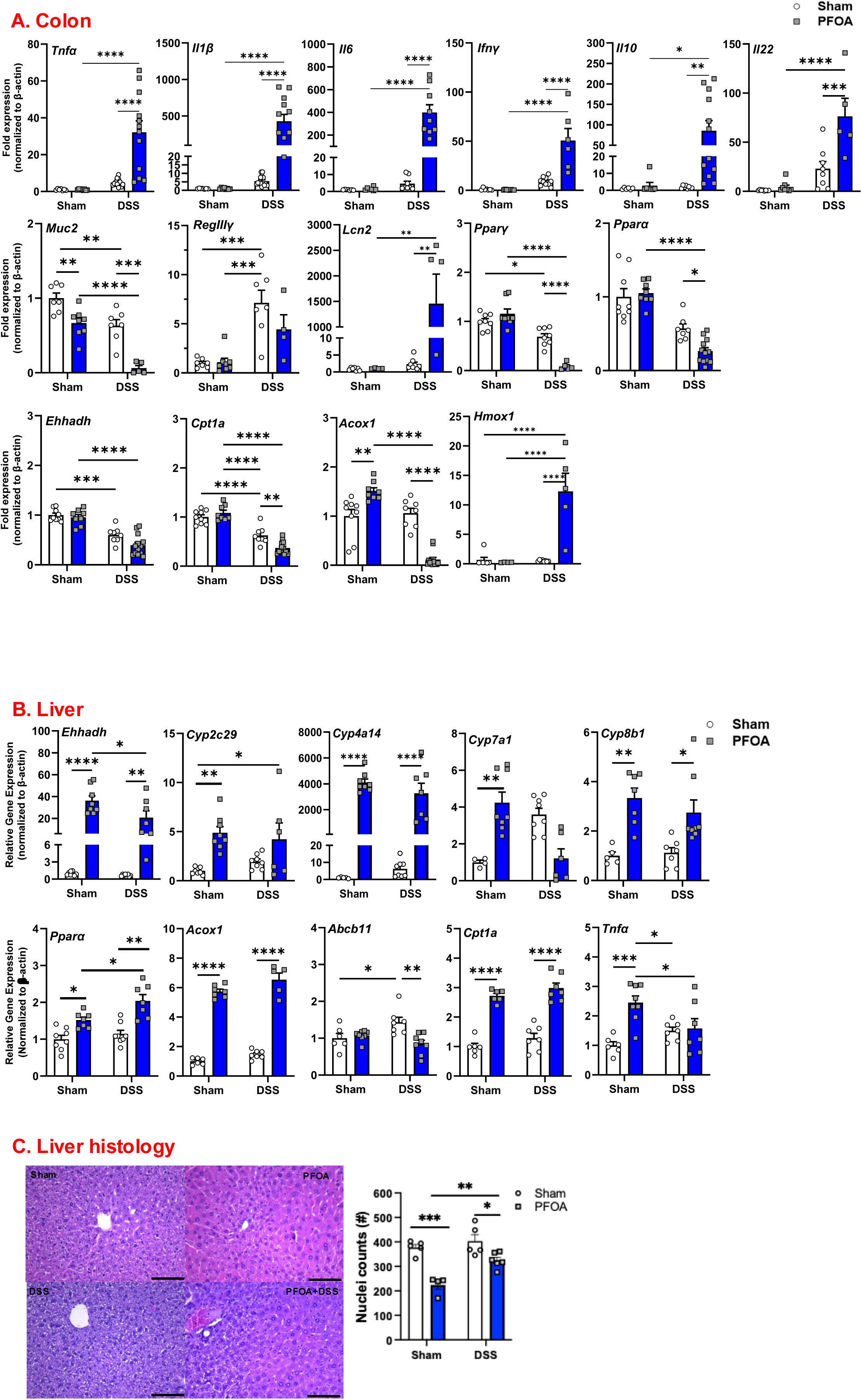
Inflammation and liver damage were exacerbated by PFOA exposure in DSS-induced colitis in a sex-dependent manner. Gene expression associated with inflammation (Tnfa, Il1b, Il6, Ifng, Il10, Il22), mucosal barrier function (Muc2, RegIIIg, Ffar2, Tlr4, Lcn2), bile acid metabolism, lipid metabolism, oxidative stress, and xenobiotic-responsive pathways (Ehhadh, Cyp2c29, Cyp4a14, Cyp7a1, Cyp8b1) was assessed by qPCR in the colon (**A**) and liver (**B**). qPCR analysis of distal colonic tissue was used to quantify cytokines regulatory and other (Ehhad, Pparg, Ppara, Acox1, Cpt1a, Hmox1) expression. (B). Representative images demonstrate exposure-associated changes in hepatic tissue architecture following induction of DSS-induced colitis (**C**). Data are presented as mean ± SEM; n=8-12, *p<0.05, **p < 0.01, ****p<0.001 by two-way ANOVA.

Prior PFOA exposure also altered metabolic and stress-response programs in the inflamed colon. DSS-induced colitis reduced expression of fatty acid oxidation and PPAR-associated genes, including *Ehhadh, Ppar*γ*, Ppar*α, and *Cpt1a*, and this reduction was further enhanced by prior PFOA exposure (**Fig 7A**). Although PFOA exposure alone increased *Acox1* expression, *Acox1* was significantly decreased during DSS-induced colitis (**Fig 7A**). In contrast, *Hmox1* expression was strongly induced in PFOA-exposed DSS-treated mice, consistent with enhanced oxidative stress or epithelial injury-associated stress responses. Thus, the male-specific worsening of DSS-induced colitis after PFOA exposure was associated with reduced epithelial barrier and metabolic adaptation, coupled with increased injury-associated inflammatory stress.

In the liver, PFOA exposure induced a distinct transcriptional program that was largely preserved during DSS-induced colitis. PFOA exposure alone increased expression of *Ehhadh, Cyp2c29, Cyp4a14, Cyp7a1*, and *Cyp8b1*, and these responses were not further enhanced by DSS-induced colitis (**Fig 7B**). PFOA exposure also increased hepatic *Ppar*α expression, which was further enhanced during DSS-induced colitis. Consistent with sustained activation of hepatic lipid metabolic pathways, the PPARα-associated fatty acid oxidation genes *Acox1* and *Cpt1a* were similarly increased in PFOA-exposed mice regardless of DSS administration (**Fig 7B**). In contrast, *Abcb11* expression was increased by DSS alone but reduced in PFOA-exposed DSS-treated mice, suggesting that prior PFOA exposure may alter bile acid transport responses during colitis (**Fig 7B**). PFOA exposure alone also increased hepatic *Tnf*α expression, although this increase was attenuated during DSS-induced colitis (**Fig 7B**). Unlike the colon, sex-dependent effects were not evident in the hepatic transcriptional response to PFOA plus DSS (**Suppl Fig 3**). Together, these findings indicate that chronic PFOA exposure drives robust hepatic xenobiotic, bile acid, and lipid metabolic transcriptional responses that are largely maintained during DSS-induced colonic inflammation, while selectively modifying bile acid transport and inflammation-associated hepatic signaling.

## DISCUSSION

As ingestion is the primary route of exposure for many environmental toxicants, the GI tract is an important target organ for exposure-associated pathology.^25^ PFAS represent one such class of chemicals, with a near-ubiquitous presence in the environment, biological persistence, and the potential to interact with epithelial cell^26–28^ and metabolic^29,30^ processes. PFAS exposure has been associated with GI disease, with elevated serum PFOA concentration reported in patients with late-onset IBD.^9^ In addition to the GI tract, the liver and immune system are major PFAS-responsive targets. In mice, oral PFOA and perfluorooctanesulfonic acid (PFOS) exposure causes hepatomegaly, altered serum markers of liver injury, disrupted lipid-associated pathways, and results in sex-dependent immune effects in mice.^10^ Here, we demonstrated that PFOA exposure beginning at weaning significantly increased GI permeability, reduced epithelial cell proliferation, altered hepatic metabolic gene expression and bile acid profiles, and reshaped the gut microbiome in mice. Lastly, PFOA exposure was also associated with increased susceptibility to DSS-induced colitis in a sex-dependent manner. Together, these findings suggest that early-life oral PFOA exposure disrupts the microbiome-gut-liver axis and may increase susceptibility to epithelial injury-induced intestinal inflammation.

The weaning period is a dynamic period of growth and maturation of the GI tract, microbiome, and immune system^31^, making it a vulnerable timeframe for environmental exposures. Oral PFOA exposure beginning at weaning increased GI permeability, inflammatory gene expression, and impaired epithelial cell dynamics compared with vehicle-exposed controls with effects that varied by tissue, dose, and exposure duration. These findings parallel a recent study in which PFOS exposure similarly produced systemic immune effects, with reduced circulating *IL4*, *IL17*α, *TNF*α, and *MCP1,* but only in male mice.^10^ Together, these findings support the concept that PFAS exposure can selectively alter immune responsiveness without causing overt inflammation.

In the liver, PFOA exposure caused significant remodeling and functional changes. This remodeling included an increased liver-to-body weight ratio, hepatocyte hypertrophy, and increased expression of metabolic and xenobiotic response genes. This is consistent with recent *in vivo* studies, in which hepatomegaly was accompanied by altered serum liver injury markers, including increased alanine aminotransferase (ALT) and alkaline phosphatase (ALP) levels and decreased triglyceride levels.^10^ These studies also demonstrate activation of Pparα-dependent pathways and lipid metabolism pathways,^10^ similar to our findings. PFOA exposure was also found to increase the expression of hepatic metabolic genes, which may affect enterohepatic bile acid circulation, xenobiotic metabolism, and systemic inflammatory tone.^32^ These findings closely align with our data and support the idea that PFOA-induced hepatic remodeling involves activation of lipid metabolic and nuclear receptor-associated pathways rather than a purely inflammatory hepatic response to exposure.

Hepatic remodeling following PFOA exposure was accompanied by alterations in circulating and hepatic bile acids. In addition to their well-characterized role in facilitating lipid digestion, bile acids act as signaling molecules that regulate metabolic, epithelial, and immune functions. After secretion into the intestine following a meal, bile acids are modified by gut microbes and reabsorbed through enterohepatic circulation. Gut microbes shape systemic and hepatic bile acid profiles and regulate enterohepatic bile acid metabolism, including hepatic Cyp7a1-dependent bile acid synthesis pathways ^33^, which were impacted by PFOA exposure. Oral PFOA exposure reduced several taurine-conjugated bile acids in both serum and liver. Taurine-conjugated bile acids can influence epithelial barrier function, microbial community structure, and inflammatory signaling.^34,35^ Therefore, PFOA-associated bile acid changes may reflect disruption of host-microbe metabolic signaling rather than isolated hepatic bile acid dysregulation. Alterations in circulating and hepatic bile acids likely reflect broader perturbations in host-microbe metabolic signaling across the liver and GI compartments. Bile acids also possess antimicrobial properties and can influence microbial community structure by selectively inhibiting or promoting the growth of specific bacterial taxa.^36^ Consistent with this idea, alpha-and beta-diversity analyses indicated that PFOA exposure altered microbial diversity and community composition.

Oral exposure to PFOA caused significant shifts in the composition of the gut microbiome. At the taxonomic level, increases in *Firmicutes*, *Actinobacteriota*, and *Proteobacteria* and reductions in *Bacteroidota* and *Verrucomicrobiota* were observed, suggesting that PFOA exposure could function as a selective environmental pressure shaping the microbial ecosystem. Many of these affected bacterial groups are known to modulate bile acid metabolism, SCFA production, and/or intestinal barrier function. For example, PFOA exposure reduced *Akkermansia* and *Lactobacillus*, taxa frequently associated with mucosal homeostasis, while expanding several *Firmicutes*- and *Proteobacteria*-associated taxa. The increase in *Escherichia/Shigella* and other *Proteobacteria*-associated taxa is also consistent with microbiome profiles often linked to inflammatory susceptibility.^37–40^ Dysregulation of bile acid metabolism has also been linked to intestinal inflammation in IBD, where dysbiosis is associated with impaired microbial bile acid transformation, increased conjugated bile acids, and reduced secondary bile acids.^41^ Although the causality of these changes cannot be determined from the current study, the parallel remodeling of bile acid and microbial composition is consistent with the disruption of host-microbe metabolic interactions within the gut-liver axis.^33,41,42^ Together with the changes in microbiome diversity and taxonomic composition, these findings indicate that oral PFOA exposure produces a broad disruption of the microbiome-gut-liver axis.

Although IBD is a multifactorial disease, the role of environmental exposures in disease severity remains poorly studied. Here, the impact of PFOA exposure on disease susceptibility and severity was assessed using genetic and epithelial injury models of colitis. DSS-induced colitis is a robust mouse model largely driven by epithelial injury and barrier disruption, followed by activation of local and recruited immune cells.^43^ Mice subjected to DSS following PFOA exposure exhibited greater disease severity than vehicle-exposed controls, with increased histological damage, barrier dysfunction, and inflammatory gene expression, but only in male mice. In contrast, PFOA did not impact disease in IL10^-/-^ mice, which may suggest that PFOA preferentially amplifies epithelial injury-driven colitis when epithelial barrier disruption is a dominant initiating event rather than a genetically driven inflammation. Consistent with this interpretation, PFOS has been reported to be immunosuppressive, suggesting that the mechanism underlying intestinal inflammation is critical to the outcome of exposure.^10^ The male-biased effect observed in our DSS model is consistent with growing evidence that PFAS-associated immune outcomes are sex-dependent.^10^ This may be due to sex-dependent immunoregulation or toxicokinetics. Interestingly, prior studies found that female mice had increased inflammation in the liver, coupled with increased circulating PFOA levels following exposure compared to male mice.^44^ In contrast, our study found PFOA-induced hepatic metabolic remodeling in both sexes; thus, sex-dependent immune regulation may be an important determinant of whether PFAS exposure translates into worsened intestinal inflammatory outcomes.

PFOA accumulation in the liver can activate PPARα, a nuclear receptor that regulates fatty acid oxidation, peroxisomal metabolism, and hepatic lipid handling.^45,46^ Although PPARα is highly expressed in hepatocytes, it is also expressed in IEC, where prior studies have implicated PPARα signaling in lipid absorption, epithelial cell programming and differentiation, and barrier function ^47^. Thus, PFOA exposure may affect GI health through both direct effects on the IEC and indirect effects resulting from altered hepatic metabolism and bile acid profiles. PPARα-regulated pathways also intersect with other nuclear receptor pathways, including the farnesoid X receptor (FXR)-dependent bile acid signaling^48^ and constitutive androstane receptor (CAR)-dependent xenobiotic responses.^49^ *Ppar*α was decreased in DSS-induced colitis in the colon, and this reduction was further enhanced by PFOA exposure. In the liver, *Ppar*α was increased by PFOA regardless of DSS-induced colitis. Together, these integrated signaling mechanisms provide a potential link between PFOA exposure, hepatic remodeling, bile acid homeostasis, microbiome, and inflammation.

Our findings identify early-life PFOA exposure as a modifier of microbiota-gut-liver homeostasis, increasing susceptibility to epithelial injury-induced colitis in a sex-dependent manner. With the persistence of PFAS in the environment and the increasing incidence of IBD worldwide, defining how these exposures alter GI and hepatic function may help identify preventable contributors to the risk of inflammatory disease. Future studies should determine whether PFOA-induced changes in colitis severity are mediated by nuclear receptor signaling, microbiota composition, or deficient epithelial repair pathways or interactions among these mechanisms.

## Acknowledgements

The authors would like to thank Dr. Sierra Schreiber for initiating some pilot studies that led to this project and Tove Slettvoll for technical assistance with the PFAS and bile acid LC/MS analysis. This project was funded by the NIH ES034806 (MG and CR), and NIH MH112507 (ASM).

## SUPPLEMENTAL FIGURE LEGENDS

**Supplemental Figure 1.**
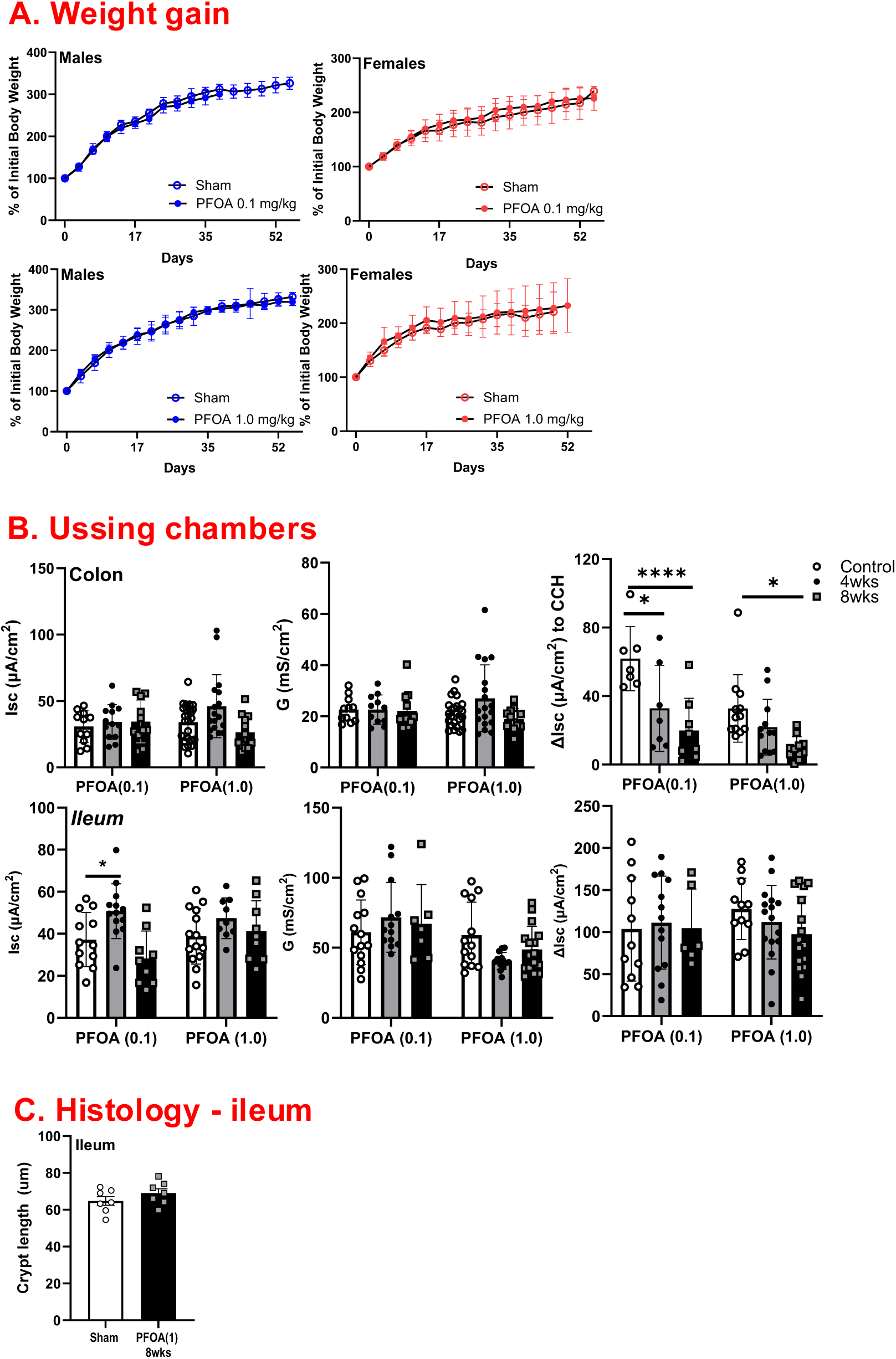
Lack of dose-dependent effects of chronic PFOA exposure on body weight, ileal histology, and intestinal physiology. Longitudinal body weight gain in male (blue) and female (red) mice following chronic exposure to varying doses of PFOA (0.1 or 1.0 mg/kg) beginning at weaning to assess dose- and sex-dependent effects on growth (**A**). Histological scoring of ileal tissue following chronic PFOA exposure (**B**). Ussing chamber analyses of ileal tissue following chronic PFOA exposure, including short-circuit current (Isc), conductance (G), and carbachol (CCH)-stimulated ion transport (**C**). Data are presented as mean ± SEM; n=8-16, *p<0.05, ****p<0.001 by ANOVA.

**Supplemental Figure 2.**
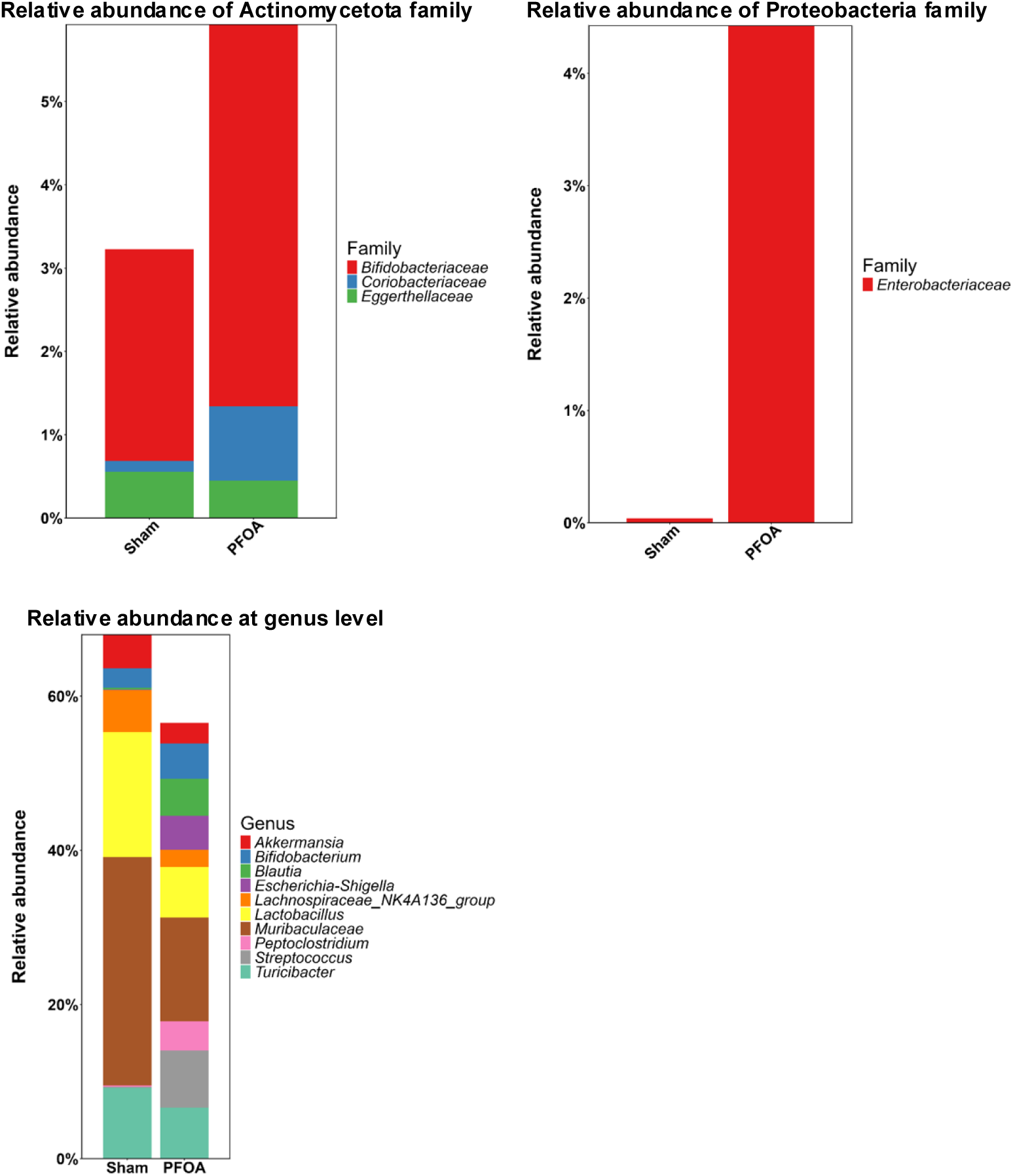
Relative abundance at the family and genus level. Relative abundance of bacterial families within the phylum Actinomycetota and Proteobacteria as well as genus-level relative abundance of fecal microbial communities in sham- vs PFOA-exposed mice. n=8 mice/group.

**Supplemental Figure 3.**
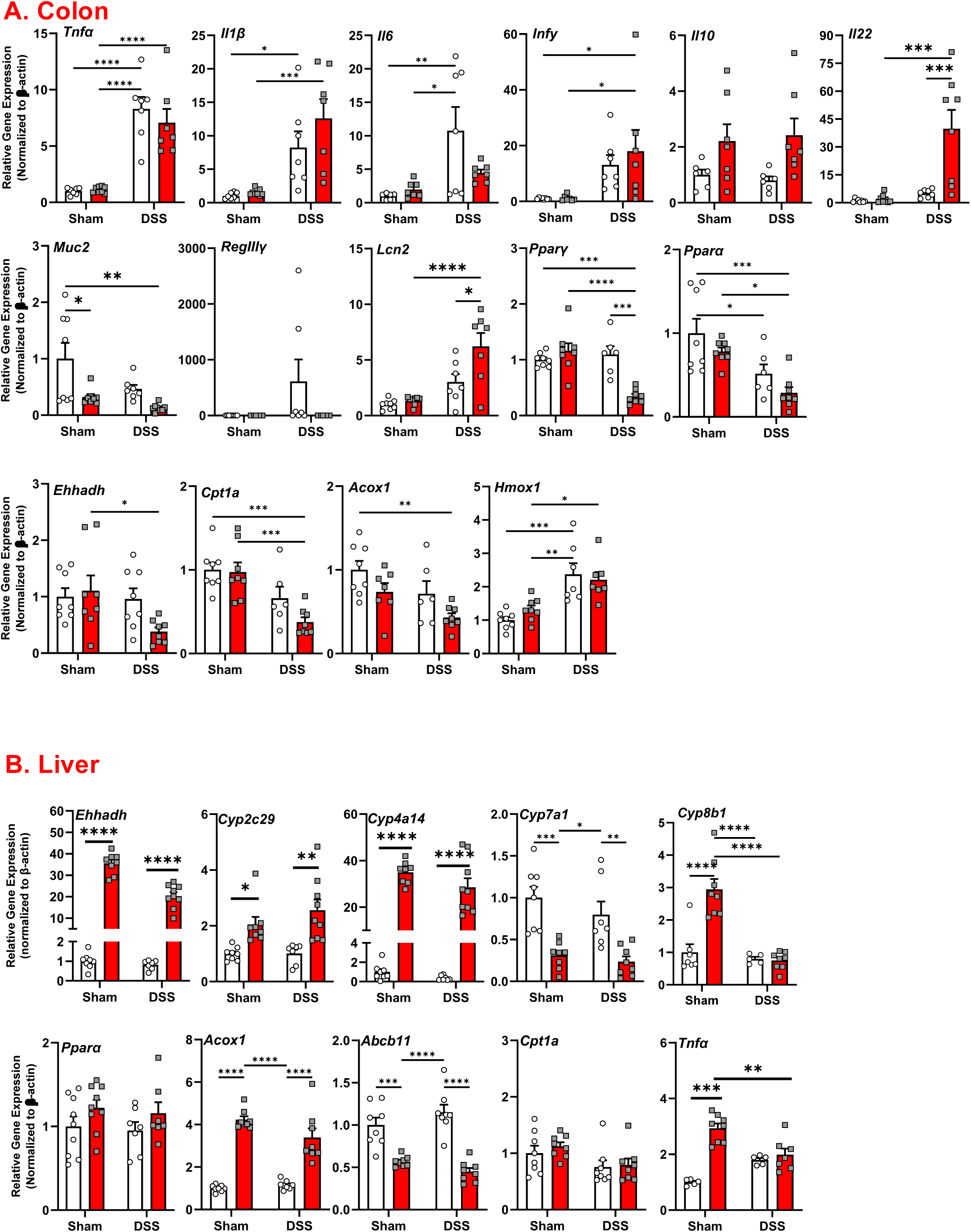
Gene expression in colon and liver in female mice exposed to PFOA + DSS-induced colitis. qPCR analysis of colon (A) and liver (B) tissue from female mice following PFOA exposure (1mg/Kg, 8 weeks) and DSS-induced colitis. Data are presented as mean ± SEM; n=8-12, *p<0.05, **p<0.01, ***p<0.001, ****p<0.0001 by two-way ANOVA.

**Supplemental Figure 4. Sex-dependent hepatic transcriptional responses following chronic PFOA exposure and DSS-induced colitis.**

qPCR analysis of liver tissue from male (**A**) and female (**B**) mice following chronic PFOA exposure and DSS-induced colitis. Data are presented as mean ± SEM; n=8-12, *p<0.05, **p<0.01, ***p<0.001, ****p<0.0001 by two-way ANOVA.

